## Supplementary material for "Epidemiological and ecological consequences of virus manipulation of host and vector in plant virus transmission": S1 Text

### Appendix S1: Comparison with previous models of vector preference

A table showing a comparison between the current model and the models of Roosien et al. (2013), Shaw et al. (2017) and Gandon (2018) is below.

| **Roosien et al. (2013)** | | **Shaw et al. (2017)** | | **Gandon (2018)** | | **Current model** | |
|---|---|---|---|---|---|---|---|
| **Description** | **Symbol** | **Description** | **Symbol** | **Description** | **Symbol** | **Description** | **Symbol** |
| *Host plant variables* | | | | | | | |
| Fraction non-infected plants | $1-I$ | Fraction healthy hosts | $H$ | Density susceptible hosts | $S$ | Density susceptible plants | $S$ |
| Fraction infected plants | $I$ | Fraction infected hosts | $I$ | Density infected hosts | $I$ | Density infected plants | $I$ |
| Plant population size = 1 | | Number of hosts/Ha<br><br>Minimum number of hosts/Ha to support vectors[1] | $F$<br><br>$\xi$ | Host density | $S+I$ | Density of plants in absence of the virus | $N$ |
| *Host plant population dynamic variables* | | | | | | | |
| Mortality rate of hosts not modelled | | | | Mortality rate of infected hosts | $d$ | Rate at which plants die naturally<br><br>Rate of roguing/disease-induced mortality | $\rho$<br><br>$\mu$ |
| *Vector variables* | | | | | | | |
| Fraction of non-inoculative vectors | $1-Z$ | Number of non-viruliferous vectors on healthy hosts/Ha<br><br>Number of non-viruliferous vectors on infected hosts/Ha | $N_h$<br><br>$N_i$ | Density of susceptible vectors | $V_s$ | Density of non-viruliferous vectors | $X$ |
| Fraction of inoculative vectors | $Z$ | Number of viruliferous vectors on healthy hosts/Ha<br><br>Number of viruliferous vectors on infected hosts/Ha | $V_h$<br><br>$V_i$ | Density of infected vectors | $V_i$ | Density of viruliferous vectors | $Z$ |
| Vector population size = 1 | | Number of vectors/Ha | $N_h + N_i + V_h + V_i$ | Density of vectors | $V_s + V_i$ | Density of vectors in the absence of the virus | $\kappa$ |

[1] This correction factor is required in Shaw et al. (2017) to "keep the model well behaved", since the model equations do not appear to include any constraint to stop the fraction of infected plants from going outside the range [0,1].

| Roosien et al. (2013) | | Shaw et al. (2017) | | Gandon (2018) | | Current model | |
|---|---|---|---|---|---|---|---|
| **Description** | **Symbol** | **Description** | **Symbol** | **Description** | **Symbol** | **Description** | **Symbol** |
| *Vector population dynamic parameters* | | | | | | | |
| Vector population dynamics not modelled | | Intrinsic vector growth rate on healthy hosts<br><br>Intrinsic vector growth rate on infected hosts | $r_h$<br><br>$r_i$ | Per capita fecundity of uninfected vectors<br><br>Per capita fecundity of infected vectors<br><br>Quality of infected hosts relative to susceptible hosts | $f_s$<br><br>$f_i$<br><br>$\phi$ | Per capita birth rate as vector population tends to 0<br><br>Proportionate change in birth rate on infected plants | $\sigma$<br><br>$\beta$ |
| | | Vector carrying capacity/healthy host<br><br>Vector carrying capacity/infected host | $K_h$<br><br>$K_i$ | Intensity of density dependence on vector fecundity<br><br>Maximal fecundities | $\kappa$<br><br>$F_s, F_i$ | Vector population density as density dependence reduces the birth rate to 0 | $\zeta$ |
| | | Mortality (and birth) rate not modelled explicitly | | Mortality rate of susceptible vectors<br><br>Mortality rate of infected vectors | $\delta_{V_s}$<br><br>$\delta_{V_i}$ | Rate at which vectors die in the absence of any effect due to number of flights<br><br>Increase in death rate as more plants visited per feed | $\alpha$<br><br>$\delta$ |
| *Transmission* | | | | | | | |
| Compound parameter: av. number of vectors per plant x freq. of vector movement x prob. that an inoculative vector transmits the virus to non-infected plant | $\phi_p$ | Maximum vector departure rate from infected plants<br><br>Rate healthy hosts become infected | $a_h$<br><br>$\beta_i$ | Probability susceptible host infected by infected vector<br><br>Probability of superinfection of an infected host<br><br>Handling time | $\beta$<br><br>$\phi_H$<br><br>$\tau$ | Average number of plants visited per viruliferous vector per day<br><br>Prob. a healthy plant inoculated by viruliferous vector in single visit | $\phi_+$<br><br>$\gamma$ |
| Compound parameter: av. number of vectors per plant x freq. of vector movement x prob. that an noninoculative vector acquires the virus from an infected plant | $\phi_v$ | Maximum vector departure rate from healthy plants<br><br>Rate vector on infected host becomes viruliferous | $a_i$<br><br>$\beta_v$ | Prob. susceptible vector gets infected on infected host<br><br>Probability of superinfection of an infected vector<br><br>Handling time | $b$<br><br>$\phi_V$<br><br>$\tau$ | Average number of plants visited per non-viruliferous vector per day<br><br>Prob. non-viruliferous vector acquires virus in single visit to infected plant | $\phi_-$<br><br>$\eta$ |

| Roosien et al. (2013) | | Shaw et al. (2017) | | Gandon (2018) | | Current model | |
|---|---|---|---|---|---|---|---|
| **Description** | **Symbol** | **Description** | **Symbol** | **Description** | **Symbol** | **Description** | **Symbol** |
| *Vector dispersal* | | | | | | | |
| Dispersal not modelled | | Dispersal loss (for each status)<br><br>Same status half-departure constant<br><br>Same status half-departure constant | $\mu$<br><br>$c_1$<br><br>$c_2$ | Dispersal not modelled | | | |
| *Vector preference* | | | | | | | |
| Preference parameter of inoculative vectors for infected plants | $\rho_+$ | Preference of viruliferous vectors for settling on infected hosts | $\epsilon$ | Searching efficiency of infected vectors on susceptible hosts<br><br>Searching efficiency of infected vectors on infected hosts<br><br>Preference for infected hosts in infected vectors (derived) | $\alpha_s$<br><br>$\alpha_i$<br><br>$\pi$ | Bias of viruliferous vectors to land on infected plants | $\nu_+$ |
| Preference parameter of noninoculative vectors for infected plants | $\rho_-$ | Preference of nonvirulent vectors for settling on infected hosts | $\delta$ | Searching efficiency of susceptible vectors on susceptible hosts<br><br>Searching efficiency of susceptible vectors on infected hosts<br><br>Preference for infected hosts in susceptible vectors (derived) | $a_s$<br><br>$a_i$<br><br>$p$ | Bias of non-viruliferous vectors to land on infected plants | $\nu_-$ |
| Cost of bias not modelled | | | | Cost associated with a bias in vector performance | $\rho$ | Cost of bias not modelled | |

<table>
<tr><th colspan="2">Roosien et al. (2013)</th><th colspan="2">Shaw et al. (2017)</th><th colspan="2">Gandon (2018)</th><th colspan="2">Current model</th></tr>
<tr><th>Description</th><th>Symbol</th><th>Description</th><th>Symbol</th><th>Description</th><th>Symbol</th><th>Description</th><th>Symbol</th></tr>
<tr><td colspan="8">Feeding Preference</td></tr>
<tr><td colspan="6" rowspan="5">Feeding not modelled</td><td>Average time spent feeding when vector chooses to settle after landing</td><td>$\Gamma$</td></tr>
<tr><td>Probability that non-viruliferous vector settles to feed on susceptible plant</td><td>$\omega_-$</td></tr>
<tr><td>Bias of non-viruliferous vector to feed on infected plant</td><td>$\epsilon_-$</td></tr>
<tr><td>Probability that viruliferous vector settles to feed on susceptible plant</td><td>$\omega_+$</td></tr>
<tr><td>Bias of viruliferous vector to feed on infected plant</td><td>$\epsilon_+$</td></tr>
<tr><td colspan="8">Recovery rates</td></tr>
<tr><td>Recovery rate of infected plants/rate of replacement of infected plants with noninfected plants</td><td>$\gamma$</td><td colspan="4">Not modelled</td><td>Replacement rate of infected plants with healthy susceptible plants</td><td>$\rho$</td></tr>
<tr><td>Recovery rate of inoculative vectors/rate of replacement of inoculative vectors with noninoculative vectors</td><td>$\tau$</td><td>Vector recovery rate by feeding[2]</td><td>$\gamma$</td><td colspan="2">Not modelled</td><td>Rate at which vectors lose infectivity</td><td>$\tau$</td></tr>
<tr><td colspan="8">Invasion criterion</td></tr>
<tr><td colspan="2">Derived</td><td colspan="2">Not derived</td><td colspan="2">Three versions derived</td><td colspan="2">Derived</td></tr>
</table>

[2] The way in which recovery of viruliferous vectors has been modelled is rather unclear from what is presented in the paper, since both model equations and associated schematic indicate that viruliferous vectors on healthy hosts "recover" to become non-viruliferous vectors on infected hosts (since the $\gamma V_h$ term links the $V_h$ and $N_i$ compartments in the model equations).
