## Supplementary material for "Epidemiological and ecological consequences of virus manipulation of host and vector in plant virus transmission": S2 Text

### Appendix S2. Deriving the Basic Reproduction Number via the Next Generation Matrix approach

Focusing on the virus-carrying compartments in Equation (26) of the main text leads to

$$\frac{dI}{dt} = \frac{\gamma Z S}{\omega_+ \Gamma(S + \nu_+ \epsilon_+ I)} - (\rho + \mu) I$$

$$\frac{dZ}{dt} = \frac{\eta \nu_- X I}{\omega_- \Gamma(S + \nu_- \epsilon_- I)} - (\tau + \alpha_+) Z$$

The disease-free equilibrium is $(S, I, X, Z) = (N, 0, \kappa, 0)$. Following the standard notation for Next Generation calculations (van den Driessche, 2017), take the rates at which new infections are caused

$$F = \begin{pmatrix} \dfrac{\gamma Z S}{\omega_+ \Gamma(S + \nu_+ \epsilon_+ I)} \\ \dfrac{\eta \nu_- X I}{\omega_- \Gamma(S + \nu_- \epsilon_- I)} \end{pmatrix}$$

and the rates at which infections are removed

$$V = \begin{pmatrix} (\rho + \mu) I \\ (\tau + \alpha_+) Z \end{pmatrix}$$

The Jacobians of these matrices are

$$J_F = \begin{pmatrix} -\dfrac{\gamma Z S \nu_+ \epsilon_+}{\omega_+ \Gamma(S + \nu_+ \epsilon_+ I)^2} & \dfrac{\gamma S}{\omega_+ \Gamma(S + \nu_+ \epsilon_+ I)} \\ \dfrac{\eta \nu_- X S}{\omega_- \Gamma(S + \nu_- \epsilon_- I)^2} & 0 \end{pmatrix}$$

and

$$J_V = \begin{pmatrix} \rho + \mu & 0 \\ 0 & \tau + \alpha_+ \end{pmatrix}$$

Evaluating $M = J_F J_V^{-1}$ at the disease-free equilibrium gives

$$M = \begin{pmatrix} 0 & \dfrac{\gamma}{\omega_+ \Gamma(\tau + \alpha_+)} \\ \dfrac{\eta \nu_- \kappa}{\omega_- \Gamma N(\rho + \mu)} & 0 \end{pmatrix}$$

The basic reproduction number, $R_0$, is then the spectral radius of the matrix $M$, i.e.

$$R_0 = \sqrt{\frac{\gamma \eta \nu_- \kappa}{\omega_- \omega_+ \Gamma^2 N(\tau + \alpha_+)(\rho + \mu)}}$$

which matches the result given in Equation (36) of the main text.
