## Supplementary material for "Epidemiological and ecological consequences of virus manipulation of host and vector in plant virus transmission": S3 Text

### Appendix S3. Finding model equilibria

#### *Disease free equilibria*

These correspond to equilibria for which $I = Z = 0$. There are two such equilibria:

- E1: $(S, I, X, Z) = (N, 0,0,0)$
- E2: $(S, I, X, Z) = (N, 0, \kappa, 0)$

where the derived parameter $\kappa$ is as defined in Equation (24), i.e.

$$\kappa = \zeta\left(1 - \frac{\alpha}{\sigma}\left(1 + \delta\left(\frac{1}{\omega_-} - 1\right)\right)\right)$$

We note that the equilibrium E1 is always biologically meaningful, whereas E2 can only be attained in practice if $\kappa > 0$.

At either equilibrium, the Jacobian of Equation (26) is given by

$$J = \begin{pmatrix} -\rho & 0 & 0 & * \\ 0 & -(\rho+\mu) & 0 & {\partial \Lambda}/{\partial \mathrm{Z}} \\ * & * & {\partial \mathrm{g}}/{\partial \mathrm{X}} - h_- & * \\ 0 & {\partial \Omega}/{\partial \mathrm{I}} & 0 & -(\tau + h_+) \end{pmatrix}$$

in which * denotes values not needed in the subsequent analysis, and where partial derivatives, and the two functions $h_-$ and $h_+$ are understood to be evaluated at the equilibrium under consideration.

The local stability of an equilibrium is controlled by the eigenvalues of $J$, i.e. by the characteristic equation $\det(J - \lambda I) = 0$. Expanding down the third column indicates that one eigenvalue is given by

$$\lambda = {\partial \mathrm{g}}/{\partial \mathrm{X}} - h_- = \sigma\left(1 - \frac{2X}{\zeta}\right) - \alpha\left(1 + \delta\left(\frac{1}{\omega_-} - 1\right)\right)$$

in which $X$ is the value at the equilibrium in question.

For E1, $X = 0$ meaning that $\lambda = \frac{\sigma\kappa}{\zeta}$ (and so that this equilibrium is definitely unstable whenever $\kappa > 0$ – i.e. whenever E2 is biologically meaningful – but potentially stable if $\kappa < 0$, depending on the other three eigenvalues).

For E2, $X = \kappa$ and so $\lambda = -\frac{\sigma\kappa}{\zeta}$ (meaning this equilibrium is potentially stable whenever $\kappa > 0$, again depending on the rest of the eigenvalues, but must definitely be unstable if $\kappa < 0$).

The remaining three eigenvalues are controlled by the eigenvalues of the 3x3 matrix

$$J^* = \begin{pmatrix} -\rho & 0 & * \\ 0 & -(\rho+\mu) & {\partial \Lambda}/{\partial \mathrm{Z}} \\ 0 & {\partial \Omega}/{\partial \mathrm{I}} & -(\tau + h_+) \end{pmatrix}$$

Expanding the characteristic equation of this second matrix down the first column indicates there is always a negative eigenvalue $\lambda = -\rho$.

The remaining pair of eigenvalues come from the 2 x 2 matrix

$$J^{**} = \begin{pmatrix} -(\rho+\mu) & {\partial\Lambda}/{\partial Z} \\ {\partial\Omega}/{\partial I} & -(\tau+h_+) \end{pmatrix}$$

This third matrix always has a negative trace, $Tr(J^{**}) = -(\rho+\mu) - (\tau+h_+)$, and has determinant

$$\Delta(J^{**}) = (\rho+\mu)(\tau+h_+) - {\partial\Omega}/{\partial I}\,{\partial\Lambda}/{\partial Z} = (\rho+\mu)(\tau+h_+) - \frac{\gamma\eta\nu_- X}{N\omega_-\omega_+\Gamma^2}$$

in which again $X$ is either $X = 0$ (E1) or $X = \kappa$ (E2).

For E1, $\Delta(J^{**}) = (\rho+\mu)(\tau+h_+) > 0$, and since $Tr(J^{**}) < 0$, both eigenvalues of $J^{**}$ are negative.

For E2, the determinant is

$$\Delta(J^{**}) = (\rho+\mu)(\tau+h_+)\left(1 - \frac{\gamma\eta\nu_-\kappa}{N\omega_-\omega_+\Gamma^2(\rho+\mu)(\tau+h_+)}\right) = (\rho+\mu)(\tau+h_+)(1-R_0^2)$$

and so if $R_0^2 < 1$ then both eigenvalues are negative, whereas if $R_0^2 > 1$ then one is positive. This means that $R_0^2 > 1$ ensures this equilibrium is unstable.

Collating the conclusions of the results above

| $\boldsymbol{\kappa > 0}$ | $\boldsymbol{R_0 > 1}$ | **Stability of E1 (Disease- and vector-free equilibrium)** | **Stability of E2 (Disease-free but vector-present equilibrium)** |
|---|---|---|---|
| ✓ | ✓ | Unstable | Unstable |
| ✓ | × | Unstable | Stable |
| × | ✓ | Stable | Not biologically meaningful |
| × | × | Stable | Not biologically meaningful |

#### *Disease present equilibria*

This corresponds to equilibria for which $I \neq 0$.

#### Case I. Vector preference does not interact with population dynamics ($\boldsymbol{\beta = 1}$, $\boldsymbol{\delta = 0}$)

Adding the equations for $dS/dt$ and $dI/dt$ from Equation (26) and equating to zero indicates

$$0 = \rho N - \rho S - (\rho+\mu)I$$

Adding the equations for $dX/dt$ and $dZ/dt$ indicates (at least when $\beta = 1$ and $\delta = 0$) that either $X + Z = 0$, which is guaranteed to generate no biologically meaningful equilibria, or that

$$X + Z = \zeta\left(1 - \frac{\alpha}{\sigma}\right) = \kappa$$

Using these expressions to eliminate $S$ and $X$ from Equation (26), then eliminating *Z,* noting throughout that $\delta = 0$ means that $h_+ = \alpha$, eventually leads to

$$I\left(\frac{\gamma\left(N - \left(1+\frac{\mu}{\rho}\right)I\right)\eta\kappa\nu_-}{\left((\tau+\alpha)\omega_-\Gamma\left(N - \left(1+\frac{\mu}{\rho}\right)I + \nu_-\epsilon_- I\right) + \eta\nu_- I\right)\left(\omega_+\Gamma\left(N - \left(1+\frac{\mu}{\rho}\right)I + \nu_+\epsilon_+ I\right)\right)} - (\rho+\mu)\right) = 0$$

Ignoring the equilibrium at $I = 0$, which has been covered above, further equilibria are given by solutions to

$$\frac{\gamma\left(N-\left(1+\frac{\mu}{\rho}\right)I\right)\eta\kappa\nu_-}{\left((\tau+\alpha)\omega_-\Gamma\left(N-\left(1+\frac{\mu}{\rho}\right)I+\nu_-\epsilon_-I\right)+\eta\nu_-I\right)\left(\omega_+\Gamma\left(N-\left(1+\frac{\mu}{\rho}\right)I+\nu_+\epsilon_+I\right)\right)}=(\rho+\mu)$$

Simple, albeit long-winded, algebraic manipulations reduce this to the quadratic

$$a_2I^2+a_1I+a_0=0$$

in which

$$a_2=\left(\nu_-\epsilon_--\left(1+\frac{\mu}{\rho}\right)+\frac{\eta\nu_-}{(\alpha+\tau)\omega_-\Gamma}\right)\left(\nu_+\epsilon_+-\left(1+\frac{\mu}{\rho}\right)\right)$$

$$a_1=N\left(\nu_-\epsilon_-+\nu_+\epsilon_++\frac{\eta\nu_-}{(\alpha+\tau)\omega_-\Gamma}+\left(1+\frac{\mu}{\rho}\right)(R_0^2-2)\right)$$

$$a_0=N^2(1-R_0^2)$$

The values of the other state variables can then be obtained by back-substitution into

$$S=N-\left(1+\frac{\mu}{\rho}\right)I$$

$$Z=\frac{\eta\nu_-\kappa I}{(\alpha+\tau)\omega_-\Gamma\left(N-\left(1+\frac{\mu}{\rho}\right)I+\nu_-\epsilon_-I\right)+\eta\nu_-I}$$

$$X=\kappa-Z$$

#### Case II. Vector preference interacts with population dynamics ($\boldsymbol{\beta}\neq\mathbf{1}$ and/or $\boldsymbol{\delta}>\mathbf{0}$)

Similar calculations indicate that again

$$S=N-\left(1+\frac{\mu}{\rho}\right)I$$

but in this case the values of $X$ and $Z$ can only be found implicitly in terms of the equilibrium value of $X$, with

$$X=\frac{(\rho+\mu)\omega_-\Gamma(S+\nu_-\epsilon_-I)((\tau+\alpha-\alpha\delta)\omega_+\Gamma(S+\nu_+\epsilon_+I)+\alpha\delta\Gamma(S+\nu_+I))}{\gamma\eta\nu_-S}$$

and

$$Z=\frac{(\rho+\mu)\omega_+\Gamma(S+\nu_+\epsilon_+I)I}{\gamma S}$$

Extensive algebraic manipulations eventually indicate that the solutions for $I$ are given by roots to the quartic

$$a_4I^4+a_3I^3+a_2I^2+a_1I+a_0=0$$

in which the coefficients are given by

$$a_0=\sigma P_0T_0-\alpha\zeta Q_0N$$

$$a_1=\sigma(P_0T_1+P_1T_0)+\alpha\zeta(Q_0L-Q_1N)$$

$$a_2=\sigma(P_0T_2+P_1T_1+P_2T_0)+\alpha\zeta(Q_1L-Q_2N)$$

$$a_3 = \sigma(P_1 T_2 + P_2 T_1) + \alpha\zeta Q_2 L$$

$$a_4 = \sigma P_2 T_2$$

and where the coefficients are defined in terms of the quantities

$$L = 1 + \frac{\mu}{\rho}$$

$$P_0 = \zeta N - C\omega_- J_0 N$$

$$P_1 = -\zeta L - C[\omega_- J_0(\nu_-\epsilon_- - L) + \omega_+ J_1 N + \eta\nu_-\omega_+ N]$$

$$P_2 = -C[\omega_- J_1(\nu_-\epsilon_- - L) + \eta\nu_-\omega_+(\nu_+\epsilon_+ - L)]$$

$$Q_0 = (1-\delta)\omega_- N J_0 + \delta N J_0$$

$$Q_1 = (1-\delta)[\omega_- N J_1 + \omega_-(\nu_-\epsilon_- - L)J_0 + \eta\nu_-\omega_+ N] + \delta[N J_1 + (\nu_- - L)J_0 + \eta\nu_- N]$$

$$Q_2 = (1-\delta)[\omega_-(\nu_-\epsilon_- - L)J_1 + \eta\nu_-\omega_+(\nu_+\epsilon_+ - L)] + \delta[(\nu_- - L)J_1 + \eta\nu_-(\nu_+ - L)]$$

$$T_0 = \omega_- N J_0$$

$$T_1 = \omega_- N J_1 + \omega_-(\nu_-\epsilon_- - L)J_0 + \eta\nu_-\omega_+ N + (\beta - 1)\nu_-\omega_-\epsilon_- J_0$$

$$T_2 = \omega_-(\nu_-\epsilon_- - L)J_1 + \eta\nu_-\omega_+(\nu_+\epsilon_+ - L) + (\beta - 1)[\nu_-\epsilon_-\omega_- J_1 + \eta\nu_-\nu_+\epsilon_+\omega_+]$$

in which

$$A = (\tau + \alpha - \alpha\delta)\omega_+\Gamma$$

$$B = \alpha\delta\Gamma$$

$$C = \frac{(\mu + \gamma)\Gamma}{\gamma\eta\nu_-}$$

$$J_0 = (A + B)N$$

$$J_1 = A\nu_+\epsilon_+ + B\nu_+ - (A + B)L$$
