## Supplementary material for "Epidemiological and ecological consequences of virus manipulation of host and vector in plant virus transmission": S4 Text

### Appendix S4. Number and stability of equilibria

We performed a randomisation scan to understand how frequently multiple biologically relevant equilibria could be obtained. We selected sets of parameters by sampling uniformly at random from the following ranges, chosen so that the full range of biologically plausible behaviours were examined.

| Parameter | Minimum value | Maximum value |
|---|---|---|
| $N$ | 1 | 2,000 |
| $\rho$ | 0.001 | 0.05 |
| $\mu$ | 0.001 | 0.2 |
| $\alpha$ | 0 | 0.6 |
| $\sigma$ | 0.001 | 0.4 |
| $\Gamma$ | 0.001 | 4 |
| $\delta$ | 0 | 1 |
| $\beta$ | 0 | 3 |
| $\tau$ | 0 | 8 (NPT); 1 (PT) |
| $\zeta$ | 1,000 (NPT); 1 (PT) | 2,400 (NPT); 300 (PT) |
| $\nu_-$ | 0.001 | 4 |
| $\nu_+$ | 0.001 | 4 |
| $\omega_-$ | 0.001 | 1 |
| $\omega_+$ | 0.001 | 1 |
| $\epsilon_-$ | 0.001 | 2 |
| $\epsilon_+$ | 0.001 | 2 |

In all cases, we calculated the values of the derived parameters $\gamma$ and $\eta$ according to the transmission type (i.e. NPT vs PT). We constrained the parameters for NPT such that $\nu_- = \nu_+$, $\omega_- = \omega_+$ and $\epsilon_- = \epsilon_+$, and – for both transmission types – immediately discarded any samples for which $\omega_\pm \epsilon_\pm > 1$.

For each of $12{,}500{,}000$ samples for each type of transmission, we then used the quartic equation in S3 Appendix to calculate the (up to) four equilibrium values for which $I \neq 0$. Solving the equations as a polynomial was computationally convenient, since we then knew in advance how many solutions to expect (i.e. four solutions, if those involving complex numbers are counted).

For certain random sets of parameters, the quartic had coefficients of vastly differing sizes (indeed, there was at least one case in which the biggest coefficient was 17 orders of magnitude larger than the smallest coefficient). This led to potential inaccuracies in the numerical solution of the quartic polynomial (for which we used the NumPy solver "roots"). To check the accuracy of the calculated roots, potential equilibria were back substituted into right hand side of the system of differential equations. Any candidate equilibria that did not lead to values sufficiently close to zero for all four equations were used as initial estimates for a direct numerical solution of the nonlinear system. In almost all cases, this two-step process found relevant solutions to sufficient accuracy. In a small number of cases the second step of the procedure either failed to identify an equilibrium or converged on a different root. These parameter sets were discarded, before replacement with valid parameter sets, thus ensuring there were $12{,}500{,}000$ samples for each type of transmission. This happened relatively infrequently: 51,472 times for NPT and 43,133 times for PT (i.e. in <0.5% of cases).

For each of the $12{,}500{,}000$ valid independent set of parameters the remained, we characterised:

- whether the vector carrying capacity $(\kappa)$ was positive or negative;
- whether the basic reproduction number $(R_0)$ was larger or smaller than one;
- the number of biologically meaningful equilibria generated by the quartic equation;
- the number of these equilibria that were also locally stable.

We defined a biologically meaningful equilibrium to be any for which the values of all state variables were non-negative at equilibrium. Stability of equilibria was determined using standard techniques based on analytic calculation of the Jacobian of Equation (26), using numerically calculated equilibrium values in the calculation of the eigenvalues of this matrix. Stability was then tested by considering the sign of the real parts.

Counts and proportions of each type of result, for each type of transmission, are tabulated below. Note that when $\kappa < 0$, it is necessarily the case that $R_0 < 1$. Also note that when rows which are in principle possible do not appear in the table, we have not proved such cases do not exist, this is simply because no such cases were found in our – very extensive – scan (an example for NPT would be when $\kappa < 0$ and $R_0 < 1$ but with two biologically meaningful equilibria which were both locally stable; we did not find a case like this in our numerical work). Finally, note that the first % column shows percentages falling into each category as a proportion of all parameter sets, whereas the second % column shows as a proportion of parameter sets for which at least one biologically meaningful equilibrium was found (i.e. after cases in which there were no biologically meaningful equilibria were filtered out).

Results are tabulated in the tables overleaf.

We identify several cases in which there are multiple stable biologically relevant equilibria:

- **Case One (shown in blue):** bistability between a locally stable disease-free equilibrium (i.e. $R_0 < 1$) at which the vector is present (i.e. $\kappa > 0$) and a single locally stable disease-present equilibrium;
- **Case Two (shown in green):** bistability between a locally stable disease-free equilibrium (i.e. $R_0 < 1$) at which the vector is absent (i.e. $\kappa < 0$) and a single locally stable disease-present equilibrium;
- **Case Three (shown in pink):** bistability between a pair of locally stable disease-present equilibria, with the disease-free equilibrium being unstable (i.e. $R_0 > 1$)
- **Case Four (shown in grey):** tristability between a locally stable disease-free equilibrium (i.e. $R_0 < 1$) at which the vector is absent (i.e. $\kappa < 0$) and a pair of locally stable disease-present equilibria
- **Case Five (shown in gold):** tristability between a locally stable disease-free equilibrium (i.e. $R_0 < 1$) at which the vector is present (i.e. $\kappa > 0$) and a pair of locally stable disease-present equilibria

Exemplars of model behaviour in Cases One and Two are already shown in the main text (e.g. Figures 5 and 9, respectively). Examples of Cases Three to Five are detailed on the following page.

Non-persistent transmission

| $\kappa > 0$ | $R_0 > 1$ | Number of biologically relevant equilibria ($I \neq 0$) | Of which how many are stable? | Number of times found in the scan | % | % (valid) |
|---|---|---|---|---|---|---|
| × | × | 0 | 0 | 9,506,452 | 76.05% | - |
| × | × | 1 | 0 | 5 | <0.01% | <0.01% |
| × | × | 1 | 1 | 17 | <0.01% | <0.01% |
| × | × | 2 | 0 | 1 | <0.01% | <0.01% |
| × | × | 2 | 1 | 486,423 | 3.89% | 21.06% |
| ✓ | × | 0 | 0 | 684,243 | 5.47% | - |
| ✓ | × | 1 | 0 | 7 | <0.01% | <0.01% |
| ✓ | × | 2 | 0 | 19,775 | 0.16% | 0.86% |
| ✓ | × | 2 | 1 | 234,040 | 1.87% | 10.13% |
| ✓ | × | 3 | 2 | 1 | <0.01% | <0.01% |
| ✓ | ✓ | 0 | 0 | 2 | <0.01% | - |
| ✓ | ✓ | 1 | 0 | 59,203 | 0.47% | 2.56% |
| ✓ | ✓ | 1 | 1 | 1,509,831 | 12.08% | 65.38% |

Persistent transmission

| $\kappa > 0$ | $R_0 > 1$ | Number of biologically relevant equilibria ($I \neq 0$) | Of which how many are stable? | Number of times found in the scan | % | % (valid) |
|---|---|---|---|---|---|---|
| × | × | 0 | 0 | 9,797,225 | 78.38% | - |
| × | × | 1 | 0 | 1 | <0.01% | <0.01% |
| × | × | 1 | 1 | 5 | <0.01% | <0.01% |
| × | × | 2 | 0 | 33 | <0.01% | <0.01% |
| × | × | 2 | 1 | 196,498 | 1.57% | 16.57% |
| × | × | 4 | 2 | 1 | <0.01% | <0.01% |
| ✓ | × | 0 | 0 | 1,516,919 | 12.14% | - |
| ✓ | × | 1 | 0 | 1 | <0.01% | <0.01% |
| ✓ | × | 1 | 1 | 1 | <0.01% | <0.01% |
| ✓ | × | 2 | 0 | 1,598 | 0.01% | 0.13% |
| ✓ | × | 2 | 1 | 127,778 | 1.02% | 10.78% |
| ✓ | × | 4 | 1 | 11 | <0.01% | <0.01% |
| ✓ | × | 4 | 2 | 11 | <0.01% | <0.01% |
| ✓ | ✓ | 1 | 0 | 3,806 | 0.03% | 0.32% |
| ✓ | ✓ | 1 | 1 | 852,607 | 6.82% | 71.90% |
| ✓ | ✓ | 3 | 1 | 96 | <0.01% | 0.01% |
| ✓ | ✓ | 3 | 2 | 3,409 | 0.03% | 0.29% |

Examples of Cases Three, Four and Five

All examples are for PT and were selected at random. The parameter sets considered are below (shown to the very large number of significant figures as generated by the randomisation procedure).

| **Parameter** | **Case Three** | **Case Four** | **Case Five** |
|---|---|---|---|
| $N$ | 1357.78471380456 | 1319.4498303372 | 758.422673812394 |
| $\rho$ | 0.0203851207824684 | 0.0405860211146373 | 0.00196785718199866 |
| $\mu$ | 0.070862116192852 | 0.0523757168662303 | 0.0423162927283986 |
| $\alpha$ | 0.364490774736621 | 0.172566453141838 | 0.0835683097305965 |
| $\sigma$ | 0.390441941465403 | 0.172671738699192 | 0.351101323665888 |
| $\Gamma$ | 0.0817319970600838 | 0.431801010900859 | 1.58321791034575 |
| $\delta$ | 0.154354048987494 | 0.266497509538162 | 0.498826196145645 |
| $\beta$ | 2.90534775065342 | 2.57881567083572 | 1.553702775786 |
| $\tau$ | 0.898696022660075 | 0.0612631271055258 | 0.0554984526167504 |
| $\zeta$ | 132.645389890262 | 145.151797209077 | 244.713013585564 |
| $\nu_-$ | 3.37903844254511 | 2.73611880606566 | 2.67039673107236 |
| $\nu_+$ | 1.96477227364072 | 0.0248878129250102 | 0.107380027200243 |
| $\omega_-$ | 0.892493274544967 | 0.966105712055771 | 0.151813191157394 |
| $\omega_+$ | 0.200137868270559 | 0.964307280496935 | 0.887821927785777 |
| $\epsilon_-$ | 0.352802530096919 | 0.474819355540881 | 0.144803724097339 |
| $\epsilon_+$ | 0.500524147329523 | 0.975518138714481 | 1.3507622398246 |

The initial conditions in the exemplar model runs shown on the next page were as follows.

| **Case** | **Initial Conditions** | $\boldsymbol{S_0}$ | $\boldsymbol{I_0}$ | $\boldsymbol{X_0}$ | $\boldsymbol{Z_0}$ |
|---|---|---|---|---|---|
| Three | 1 | 1000 | 1 | 100 | 1 |
| | 2 | 8.8 | 0.4 | 0.34 | 0.17 |
| Four | 1 | 5 | 550 | 8 | 50 |
| | 2 | 600 | 300 | 3 | 7 |
| | 3 | 1300 | 1 | 0.001 | 0.001 |
| Five | 1 | 5 | 100 | 8 | 50 |
| | 2 | 20 | 150 | 4 | 1 |
| | 3 | 760 | 0.5 | 20 | 0.5 |

Graphs showing examples of Cases Three, Four and Five

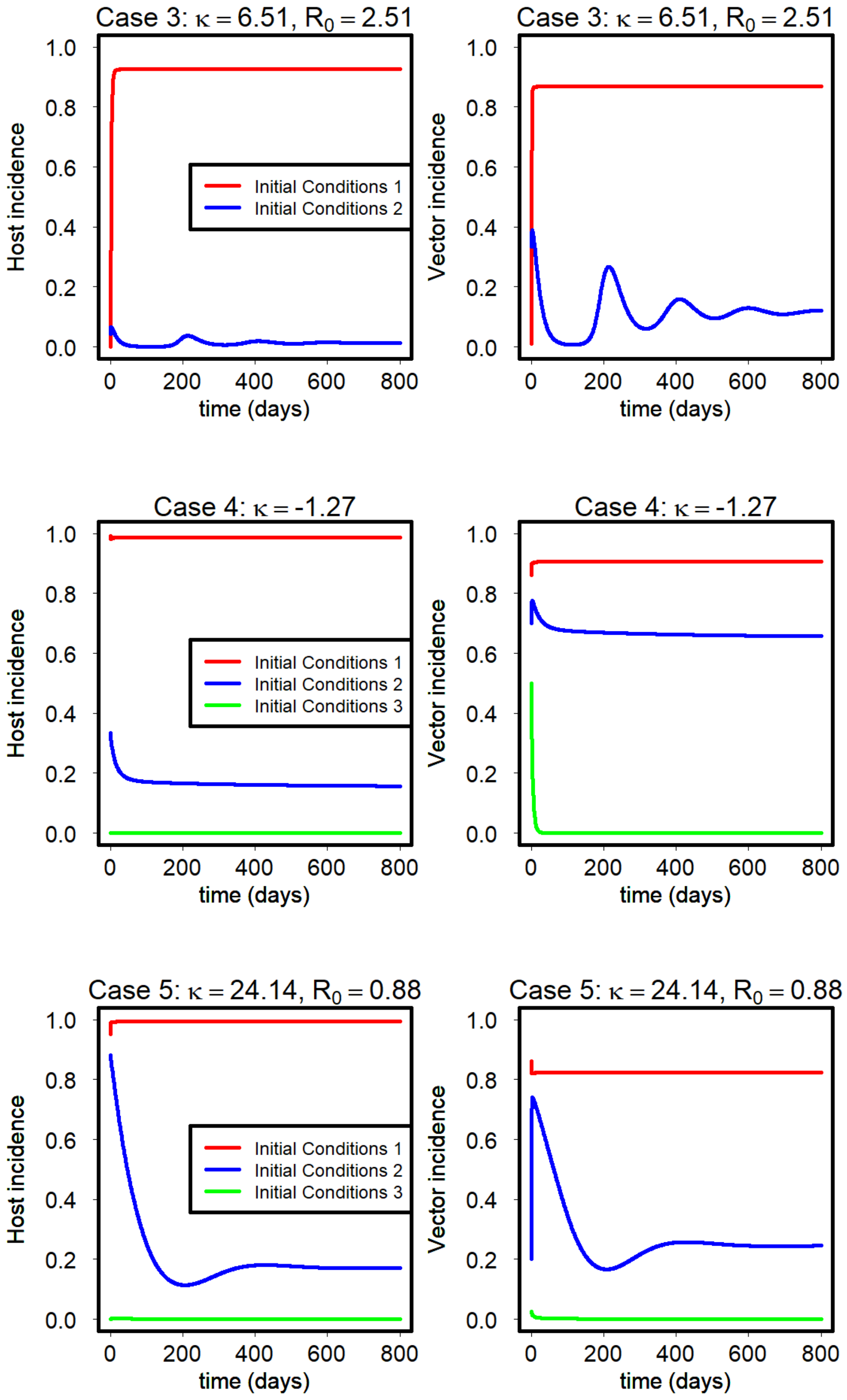
